# Differential requirement of m^6^A reader proteins, IGF2BP2 and HNRNPA2B1 for the processing of N^6^-methyladenosine modified H19 lncRNA: Stability versus miR-675 biogenesis

**DOI:** 10.1101/2024.08.07.606971

**Authors:** Samarjit Jana, Abhishek Chowdhury, Kumaravel Somasundaram

## Abstract

H19, a lnc-pri-miRNA that encodes miR-675, is dysregulated in numerous cancers. However, the specific mechanisms underlying H19 processing, particularly miR-675 formation, remain unclear. Our study reveals that H19 is highly expressed and m^6^A modified in a METTL3-dependent manner in glioblastoma (GBM) and glioma stem cells (GSCs). Silencing METTL3 reduced both H19 and miR-675 levels, whereas overexpressing METTL3 promoted miR-675 processing without affecting H19 levels. Further, miR-675 derived from exogenously expressed H19 was affected considerably more in METTL3 silenced glioma cells compared to H19 levels, suggesting differential requirements in the processing of m^6^A modified H19 transcript. We demonstrate that H19 interacts with m^6^A reader proteins, IGF2BP2 and HNRNPA2B1, and silencing either reduced H19 and miR-675 levels. However, a high level of miR-675 seen in METTL3 overexpressing cells is severely affected in HNRNPA2B1-silenced compared to IGF2BP2-silenced glioma cells. Interestingly, IGF2BP2 silencing more significantly affected H19 stability from exogenous H19 construct, while HNRNPA2B1 silencing severely impacted miR-675 processing. Site-directed mutagenesis confirmed the presence of two m^6^A sites in the first exon of H19, with site #1 facilitating HNRNPA2B1 interaction to promote miR-675 processing. In contrast, the IGF2BP2 interaction is promoted by site #2, resulting in enhanced H19 stability. H19-METTL3-HNRNPA2B1-miR675 axis inhibited Calneuron 1 (CALN1), a known target of miR-675, to promote glioma cell migration. Notably, a low CALN1/high H19 predicted a poor prognosis in GBM patients and was further exacerbated by a high METTL3 or HNRNPA2B1 but not IGF2BP2 transcript levels. Thus, we found that the H19 transcript is highly expressed in GBM and m^6^A modified, and the m^6^A reader proteins, IGF2BP2 and HNRNPA2B1, regulate the H19 processing differently to promote glioma cell migration.

## Introduction

Long non-coding RNAs (lncRNAs) have emerged as critical regulators of various cellular processes, including gene expression, chromatin remodeling, and cellular signaling (Fazi et al., 2018; Keniry et al., 2012; J. Wang et al., 2019; W. Wu et al., 2017; Xu et al., 2021). Among these mechanisms, the interaction between lncRNAs and microRNAs (miRNAs), including lncRNA-derived primary miRNA transcripts (lnc-pri-miRNA), has garnered significant interest in recent years (D. He et al., 2021). Recent studies have highlighted the importance of dynamic RNA modifications in modulating lncRNA function and stability. Among these modifications, N^6^-methyladenosine (m^6^A) is the most prevalent and well-studied internal modification in eukaryotic mRNA and lncRNA transcripts (R. Z. He et al., 2020a; Visvanathan et al., 2017, 2019b; Yue et al., 2015). LncRNAs exhibit dynamic m^6^A modification patterns that influence their stability, localization, and interactions with other RNA-binding proteins (R. Z. He et al., 2019, 2020b).

The H19 gene, located on chromosome 11p15.5, encodes a maternally expressed long non-coding RNA that has been implicated in diverse cellular processes, including proliferation, differentiation, and apoptosis (Shermane Lim et al., 2021). The H19/miR-675 axis, comprising the long non-coding RNA H19 and its processed microRNA, miR-675, has emerged as a complex regulatory network with profound implications for cancer development and progression (Cai & Cullen, 2007). This axis exhibits highly disparate expression patterns across various cancer types, underscoring its multifaceted role as both a promoter and suppressor of tumor progression. In this context, the H19/miR-675 axis functions as a double-edged sword, exerting context-dependent effects on cancer development and metastasis (Shermane Lim et al., 2021). H19 has been shown to promote cancer cell proliferation, invasion, metastasis, and resistance to chemotherapy (Guan et al., 2016; L. Liu et al., 2016; Ma et al., 2018; Matouk et al., 2014; J. Wang et al., 2019; Zhang et al., 2020). Other reports propose tumor suppressor functions for H19 (Y. Hao et al., 1993; Juan et al., 2000).

Glioblastoma (GBM) is the most common and aggressive adult brain tumor with a dismal prognosis (Stupp et al., 2009). Glioma stem-like cells (GSCs) are the main culprits behind highly aggressive GBM progression (Visvanathan et al., 2017). They also cause tumor relapse after remission because of their radio- and chemoresistant nature. High levels of H19 seen in GBM are shown to promote proliferation, invasion, angiogenesis, and stemness of GBM cells (Jiang et al., 2016; W. Li et al., 2016; Shi et al., 2014; W. Wu et al., 2017).

The H19/miR-675 axis is shown to have a dichotomous expression pattern in a context-dependent manner. H19 and miR-675 derived from H19 show a positive correlation in their expression pattern (L. Liu et al., 2016; Shi et al., 2014). However, a negative correlation between H19 and miR-675 is reported in pancreatic ductal carcinoma (Ma et al., 2018). Regulation of H19 and miR-675 expression by m^6^A modification has been proposed (J. Hao et al., 2020). Thus, the diverse expression patterns of H19 and miR-675 appear to contribute to the divergent role in cancer progression despite their origin from the same genomic locus and sharing a common transcriptional regulatory sequence. Understanding the molecular mechanisms underlying the dysregulation of this axis in cancer may provide valuable insights into the development of novel therapeutic targets for cancer treatment. In this study, we identified that H19 is highly expressed in GBM and GSCs compared to normal brain samples and DGCs (differentiated glioma cells). We also found that H19 undergoes METTL3-dependent m**^6^**A modification. More importantly, we demonstrate that two m^6^A reader proteins, HNRNPA2B1 and IGF2BP2, play a differential role in H19 stability versus its processing to form miR-675. Thus, we have discovered the complex mechanism behind H19/miR-675 expression involving m^6^A modification and its reader proteins.

## Materials and Methods

### Cell lines

The U87, U343, U252, LN229, A172, and U373 glioblastoma cell lines were obtained from ATCC, and the immortalized human astrocyte cell line IHA (NHA-hTERT-E6/E7) was acquired from Dr. Russell Pieper’s laboratory at the University of California, San Francisco (San Francisco, CA). These cell lines were cultured in specific media under standard conditions, maintaining a 5% CO2 atmosphere in a humidified incubator at 37°C. Mycoplasma contamination was assessed using RT–PCR, and all cell lines were confirmed to be free of mycoplasma using the EZdetect PCR Kit for Mycoplasma Detection (HiMedia).

### Plasmids and siRNAs

The shRNA plasmid (pLKO.1) for METTL3 (TRCN0000289814, TRCN0000289743, TRCN0000289812, TRCN0000289813, TRCN0000289742), IGF2BP2 (TRCN0000149002, TRCN0000149224, TRCN0000146301, TRCN0000148718, TRCN0000148565), and HNRNPA2B1 (TRCN0000010582) were obtained from the TRC library (Sigma, IISc). The over-expression constructs of METTL3 were obtained from Origene (#RC200869, U.S.A), and the over-expression constructs of H19 were obtained from Addgene (#122473, U.S.A). 4s1m-H19 and 4s1m-control plasmid obtained from Prof. Dung-Fang Lee’s lab (University of Texas, USA). Control siRNA and siRNA used against METTL3 were purchased from GE HealthCare Dharmacon Inc (On-TARGET plus Human siRNA SMART pool). The hsa-miR-675-5p *mir*Vana® miRNA mimic (#4464066) and Negative control (#4464058) were purchased from Thermo Fisher Scientific.

### Antibodies and reagents

Primary antibodies were purchased from the following commercial vendors: anti-m^6^A (#D9D9W) was purchased from Cell Signaling Technology, anti-METTL3 (#ab195352), anti-hnRNPA2B1 (#ab183654), and anti-IGF2BP2 (#ab128175) from Abcam, anti-GAPDH, and anti-β-actin from Sigma Aldrich. Goat anti-mouse HRP conjugate was purchased from Bio-Rad (#170-5047, WB 1:5000), goat anti-rabbit (H+L) secondary HRP conjugate was purchased from Invitrogen (#31460, WB 1:5000), Actinomycin D was purchased from Sigma Aldrich (#A1410-2MG).

### Bioinformatic analysis

Normalized gene expression profiles for GBM and control samples and associated clinical information were acquired from multiple RNA-Seq and microarray datasets through the GlioVis data portal (http://gliovis.bioinfo.cnio.es/). Gene-level read counts and clinical data for GBM and control samples in the CGGA RNA-Seq dataset were retrieved from the Chinese Glioma Genome Atlas (CGGA) database (http://www.cgga.org.cn/), normalized to effective library size and variance stabilized with DESeq2 (version 1.43.5) (Love et al., 2014). Normalized expression data of genes in GSC and DGC samples was obtained from Suvà et al. (GSE54792) (Suvà et al., 2014b). Gene-level read count data were obtained for low-grade glioma patients from TCGA LGG RNA-Seq and miRNA-Seq datasets using TCGAbiolinks R package (version 2.18.0) (Colaprico et al., 2016) and subsequently normalized with DESeq2. TPM-normalized gene expression data from control tissues across different origins were sourced from the Genotype-Tissue Expression (GTEx) Portal (https://gtexportal.org/home/). Gene expression data normalized by RSEM were acquired for 31 distinct cancer types from the TCGA Pan-Cancer Atlas Project through the TCGA Expression Browser (https://tools.altiusinstitute.org/tcga/), and cancers lacking normal samples were excluded from the analysis.

FASTQ files corresponding to RNA-Seq and RIP-Seq samples for shNT and shMETTL3 were retrieved from the National Center for Biotechnology Information (NCBI) Sequence Read Archive (SRA) under accession number SRP163326 using the SRA Toolkit. To ensure data quality, raw sequencing reads were initially subjected to quality assessment using FastQC (version 0.12.1; http://www.bioinformatics.babraham.ac.uk/projects/fastqc/). The latest human reference genome assembly (GRCh38) and corresponding gene annotation file in GTF format were retrieved from the GENCODE project (Release 45) (https://www.gencodegenes.org/) for use in downstream analysis. Reads were aligned to the human reference genome (GRCh38) using STAR (version 2.7.11b) (Dobin et al., 2013). Gene-level quantification of RNA-seq reads was performed using featureCounts (version 2.0.6) (Liao et al., 2014). Differential expression analysis was carried out using DESeq2, wherein raw read counts were normalized using estimated size factors, and differential expression was evaluated using a negative binomial generalized linear model (GLM) with shrinkage estimation of dispersions and fold changes. Genes were considered differentially expressed at │log_2_FC│ ≥ 0.58, and Benjamini-Hochberg adjusted p-value < 0.05. Aligned RIP-Seq BAM files were subjected to filtering using Sambamba (version 1.0.1) (Tarasov et al., 2015) to exclude unmapped and multi-mapped reads. Filtered BAM files were processed using BEDTools (version 2.31.1) (Quinlan & Hall, 2010) to remove reads aligning to blacklist regions. Peaks were identified using the protocol described by Visvanathan et al. (Visvanathan et al., 2019a). Genomic tracks depicting read coverage and signal enrichment were visualized using the Gviz R package (version 1.48.0) (Hahne & Ivanek, 2016).

Genes shared between TCGA GBM (RNA-Seq) and GSC-DGC cohorts were identified and those with │log_2_FC│ ≥ 0.58 and p-value < 0.05 were considered differentially regulated. Differential enrichment of m^6^A peaks identified in shNT and shMETTL3 conditions was calculated by subtracting the fold enrichment in shNT from that in shMETTL3. The screening thresholds for differentially m^6^A-modified genes were set at │log_2_FE│ ≥ 0.58 and p-value < 0.05.

A comprehensive non-redundant list of RNA binding proteins (RBPs) was curated by integrating data from Bhargava et al. (Bhargava et al., 2017), EuRBPDB (J. Y. Liao et al., 2020), RBPbase (https://apps.embl.de/rbpbase/), and RBPDB (Cook et al., 2011). Pearson’s correlation analysis was carried out to identify RBPs positively correlated with H19 in the TCGA GBM (RNA-Seq) cohort. Experimentally validated RBPs binding to H19 in human and mouse were curated from ENCORI (J. H. Li et al., 2014). The intersection of these three gene sets identified experimentally validated RBPs conserved in both human and mouse that exhibit a positive correlation with H19.

To assess the prognostic significance of gene expression levels, patients were stratified into high and low gene expression groups based on optimal cutoff values for gene expression determined using the *surv_cutpoint* function in the survminer R package (version 0.4.9). Cox proportional hazards regression incorporated in the survival R package (version 3.6.4) was used to estimate hazard ratios (HRs) and evaluate the impact of gene expression levels on overall survival. The log-rank test was applied to assess significant differences in survival probabilities between patient subgroups. Forest plot summarizing survival outcomes was generated using the forestplot R package (version 3.1.3).

### RNA interference and lentivirus preparation

METTL3 knockdown was achieved by transfecting METTL3 siRNA using Dharmafect1 transfection reagent in Opti-MEM (Invitrogen, #22600-050) medium. Cells were harvested at 24-, 48-, 72-, and 96-hours post-transfection to assess knockdown efficiency.

For Lentivirus preparation of shRNA (pLKO.1 vector), HEK-293T cells were transfected with shRNA plasmids and helper plasmids psPAX2 and pMD2.G using Lipofectamine 2000 (Invitrogen #11668027) in Opti-MEM (Invitrogen #22600-050) medium. After 6 hours, the Opti-MEM medium was replaced with fresh DMEM supplemented with 10% FBS, and the virus was collected after 60 hours of transfection. METTL3, IGF2BP2, and HNRNPA2B1 knockdowns were accomplished by infecting cells with shMETTL3, shIGF2BP2, and shHNRNPA2B1 lentivirus, respectively, followed by puromycin selection.

### Virus packaging 4s1m-H19 and infection

The TetO-FUW-4S1m or TetO-FUW-4S1m-H19 plasmids were co-transfected with packaging plasmids psPAX2 and pMD2.G into HEK-293T cells. The resulting virus was harvested from the cell culture medium after 48 hours. The glioma cell line LN229 was then infected with these viral particles along with the M2rtTA virus, which was similarly produced in HEK-293T cells. The infection was carried out in the presence of 8ug/ml polybrene (Sigma-Aldrich, USA). Thirty-six hours post-infection, HEK293T and LN229 cells were treated with 1μg/ml doxycycline (Sigma-Aldrich, USA) for 24 hours to induce the expression of 4S1m or 4S1m-H19.

### Western blot analysis

Total proteins were isolated using RIPA buffer supplemented with protease and phosphatase inhibitors (Sigma, USA). Protein concentrations were determined with Bradford’s reagent, utilizing a standard BSA curve. Equal protein amounts from all conditions were mixed with 1x protein loading dye, denatured at 95°C for 15 minutes, loaded, and separated by SDS-PAGE. Proteins were then transferred to PVDF membranes (Merck Millipore, Billerica, MA, USA), which were blocked in 5% skim milk for 1 hour at room temperature. The membranes were incubated with primary antibodies overnight at 4°C. After three washes with TBST, the PVDF membranes were incubated with HRP-conjugated goat anti-mouse or anti-rabbit secondary antibodies for 1 hour at room temperature. Finally, the blots were washed and developed using the GE ImageQuant machine with Bio-Rad Clarity and Clarity Plus ECL chemiluminescent reagent.

### RNA isolation and qRT-PCR analysis

Total RNA was extracted using TRI reagent (Sigma, U.S.A.), and 2µg of RNA underwent reverse transcription using the High-capacity cDNA reverse transcription kit (Life Technologies, USA) following the manufacturer’s instructions. Subsequently, qRT-PCR was conducted using the DyNAmo ColorFlash SYBR Green qPCR Kit on the ABI Quant Studio 5 Sequence Detection System (Life Technologies, USA). Gene expression analysis was performed using the ΔΔCt method with ATP5G and GAPDH as internal control genes. Real-time primer details are provided in a table.

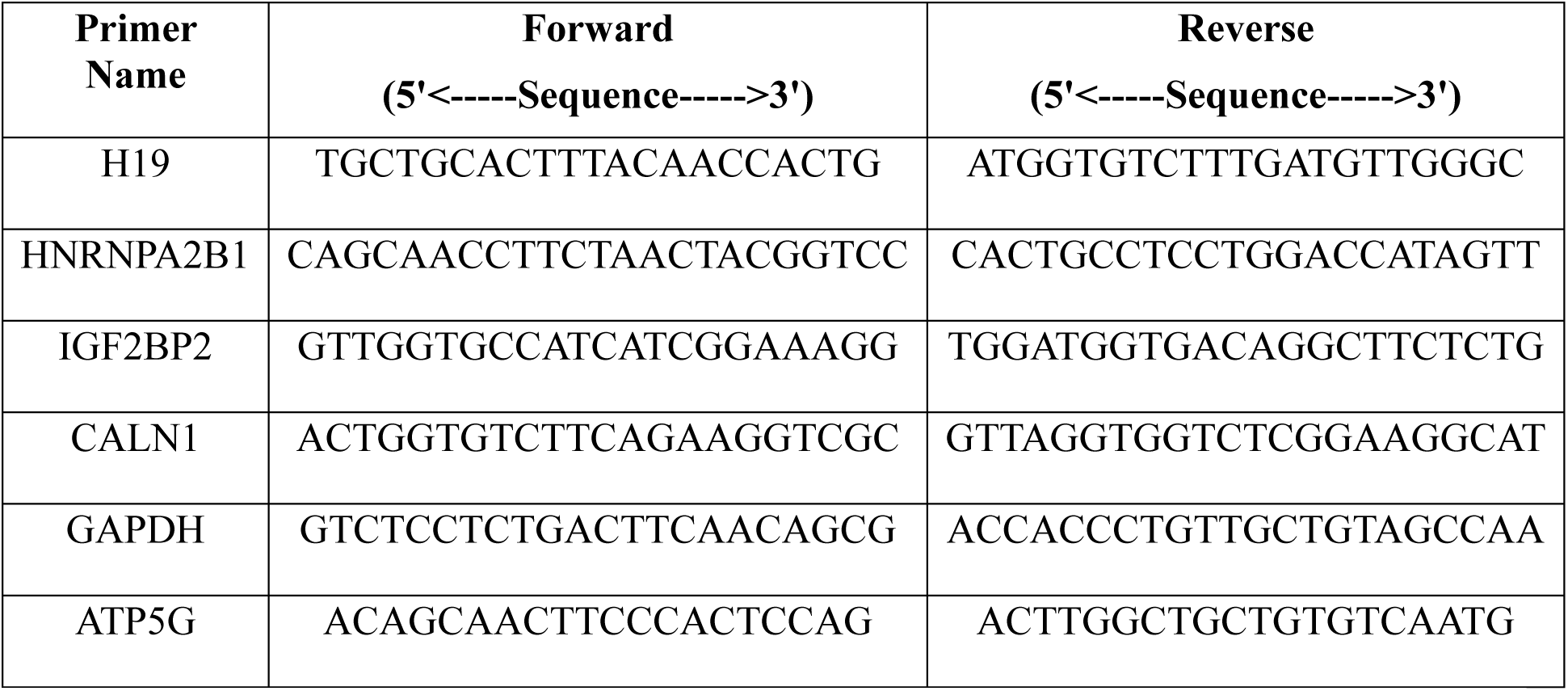

### miRNA isolation and quantification

The total RNA was isolated from LN229 cells using a miRNeasy kit (Qiagen, Germany, #217004). Isolated RNA samples were normalized to 10 ng of input for reverse transcription and cDNA synthesis using TaqMan Advanced miRNA cDNA Synthesis Kit (Invitrogen, ThermoFisher Applied Biosystems™, #A28007) according to the manufacturer’s protocol. Each cDNA sample (5 µL) was added to reaction plates containing 15 µL of TaqMan Fast Advanced Master Mix (ThermoFisher Applied Biosystems™, #4444556). The qRT-PCR for has-miR-675-5p (# A25576) and has-miR-191-5p (# A25576) were performed on Applied Biosystems Quant Studio 5 thermocycler with 1 cycle of 95℃ for 20 sec, followed by 40 cycles of 95℃ for 1-sec denaturation and 60℃ for 20 sec (annealing/extension) and normalized by the miR-191-5p according to the manufacturer’s guidelines.

### RNA stability assays

LN229 cells were exposed to 5 μg/ml actinomycin D for different durations as specified (0, 2, 4, and 6 hours). Following treatment, RNA was extracted, cDNA was synthesized, and qRT– PCR was conducted as previously outlined. The RNA half-life (t_1/2_) was determined through linear regression analysis.

### Mutagenesis

For H19 mut#1-4s1m and H19 mut#2-4s1m A-G point mutation, the sites were mutated using the Q5 Site-Directed Mutagenesis kit (#E0554S, NEB) according to the manufacturer’s instructions. Mutations were confirmed by Sanger sequencing. Mutation primers are provided as a table.

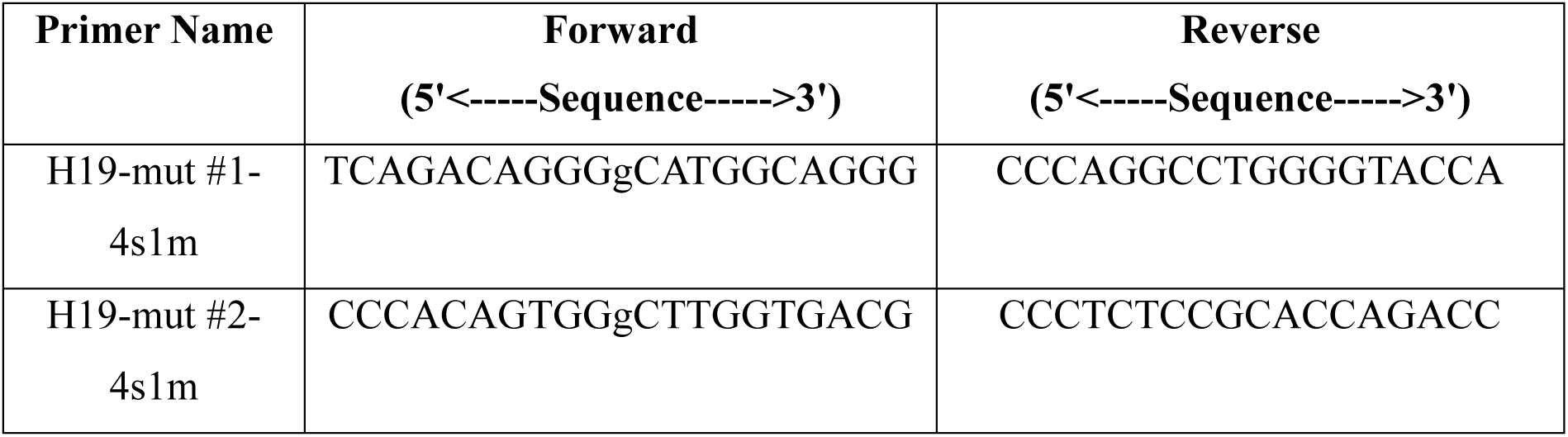

### RNA immunoprecipitation (RIP) assay

RIP assays were performed as described earlier (Visvanathan et al., 2017). Briefly, cell lysate was incubated with METTL3, IGF2BP2, and HNRNPA2B1 antibodies-coated magnetic beads at 4 °C for 24 h. The protein-associated RNAs were eluted and purified for further qRT-PCR analysis.

### m^6^A RIP-PCR assay

The rabbit anti-m^6^A antibody (#E1610S, NEB, U.S.A) was conjugated to Protein A agarose beads (Sigma, U.S.A) in 200 µl of 1 M IP Buffer (1 M NaCl, 10 mM sodium phosphate, 0.05% Triton-X) for 4 hours at 4°C. Subsequently, the beads underwent three washes in 200 µl of 140 mM IP Buffer (140 mM NaCl, 10 mM sodium phosphate, 0.05% Triton-X). RNA was denatured by incubating at 75°C for 5 minutes, then by cooling on ice for 2-3 minutes and binding to antibody-conjugated beads in 200 µl of 140 mM IP Buffer overnight at 4°C. The beads were then treated with 100 µl Elution Buffer (5 mM Tris-HCL pH 7.5, 1 mM EDTA pH 8.0, 0.05% SDS, 4.2 µl Proteinase K (20 mg/ml)) for 2 hours at 50°C, and RNA was isolated using phenol:chloroform extraction followed by ethanol precipitation.

### Purification of the H19-interacting protein complex

H19-interacting proteins were isolated using streptavidin-binding aptamers S1m, following previously described methods (Leppek & Stoecklin, 2014). Briefly, LN229 cells expressing 4S1m-H19 were suspended in 500 μl ice-cold lysis buffer (containing 20 mM Tris-HCl pH 7.5, 150 mM NaCl, 1.5 mM MgCl2, 2 mM DTT, 2 mM RNase inhibitor, and a complete protease inhibitor cocktail). The lysates were then centrifuged for 5 minutes at 12,000 rpm at 4°C. The resulting supernatants were incubated overnight at 4°C with streptavidin agarose beads (Thermo Fisher Scientific, USA) under rotation. Subsequently, the streptavidin agarose beads were washed five times for 5 minutes at 4°C with wash buffer (containing 20 mM Tris-HCl pH 7.5, 300 mM NaCl, 5 mM MgCl2, and 2 mM DTT). Finally, the protein complexes associated with 4S1m-H19 were subjected to SDS-PAGE electrophoresis.

### Transwell migration and invasion assay

After transfection, 1 × 10^5^ cells were suspended in a serum-free medium and placed in the upper chambers of transwell filters without Matrigel (migration assay) or coated with Matrigel (invasion assay) (Corning, Merck, USA). The lower chamber was filled with a complete growth medium containing FBS. The cells were incubated at 37°C for 12 hours for the migration assay and 48 hours for the invasion assay in a 5% CO2 atmosphere. Subsequently, cells on the upper side of the transwell membrane were removed using cotton swabs. The migrated cells were fixed with methanol and stained with 0.1% crystal violet. They were then counted in five randomly selected fields per well using a light microscope (Olympus, Tokyo, Japan).

### Wound healing assay

Glioma cells (LN229) were seeded into six-well culture plates and grown until they reached 80-90% confluence. Subsequently, a straight-line scratch was made in the cell layer using sterile 100-μl pipette tips, and the media was replaced with serum-free media. Images were captured using a microscope 12 hours after the scratches were made, and the distance of wound healing was measured and compared to the initial time point (0 hours).

### Statistical analysis

Statistical analyses were conducted using either GraphPad Prism 6 (GraphPad Software, La Jolla, CA) or R. Unpaired t-tests or one-way ANOVA, followed by Tukey’s test for post-hoc pairwise comparisons, were employed as appropriate for each experiment. Data are expressed as means ± s.e.m. or means ± s.d. All experiments were performed in triplicate unless otherwise stated. A significance level of p < 0.05 was deemed statistically significant.

## Results

### N^6^-methyladenosine modification regulates the stability of lncRNA H19 differently from its processing to miR-675

To identify clinically relevant lncRNAs concerning GBM, we intersected glioblastoma (GBM)-regulated (Paul et al., 2018) with Glioma stem cell (GSC)-regulated transcripts (Suvà et al., 2014a), which resulted in the identification of several regulated mRNAs and lncRNAs (**Figure 1A**; **Supplementary Table 1**). An overlap of this set of transcripts with m^6^A modified (m^6^A RIP-Seq) and METTL3 regulated transcripts (RNA-Seq) in GSCs (Visvanathan et al., 2019); **Figure 1B**; **Supplementary Table 2**) identified several transcripts that are GBM and GSC upregulated as well as m^6^A modified in a METTL3-dependent manner (**Figure 1C**; **Supplementary Table 3, 4, 5, 6 and 7**). Among them, there were only two lncRNAs, H19 and FAM95B1 (**Figure 1C)**. We have recently demonstrated that FAM95B1 is a **p**53-**i**nactivating **T**RIM28-**a**ssociated **l**ncRNA (PITAR) (Jana et al., 2023). H19 lncRNA, which is the host lncRNA for miR-675, was chosen for further studies for various reasons. H19 is upregulated in GBM (log_2_FC: 4.80) and GSC (log_2_FC: 3.21), m^6^A modified in a METTL3-dependent manner (Peak enrichment score: 2.17) and regulated by METTL3 (log_2_FC: 7.18) (**Figure 1D**, **E, and F**). H19 was the first lncRNA to be described. It is 2.36-kilobase long with five exons (**Figure 2A**). The first exon codes for the miR-675 (Cai & Cullen, 2007). While conflicting roles in tumorigenesis for H19 have been reported, most reports indicate an oncogenic function (Shermane Lim et al., 2021). H19 and miR-675 derived from it show a higher level of expression in GBM compared to lower-grade glioma (**Supplementary Figure 1A and B**), with a significant positive correlation among them (**Supplementary Figure 1C**). H19 and miR-675 show a varying expression pattern among glioma cell lines (**Supplementary Figure 1D**).

**Figure 1:**
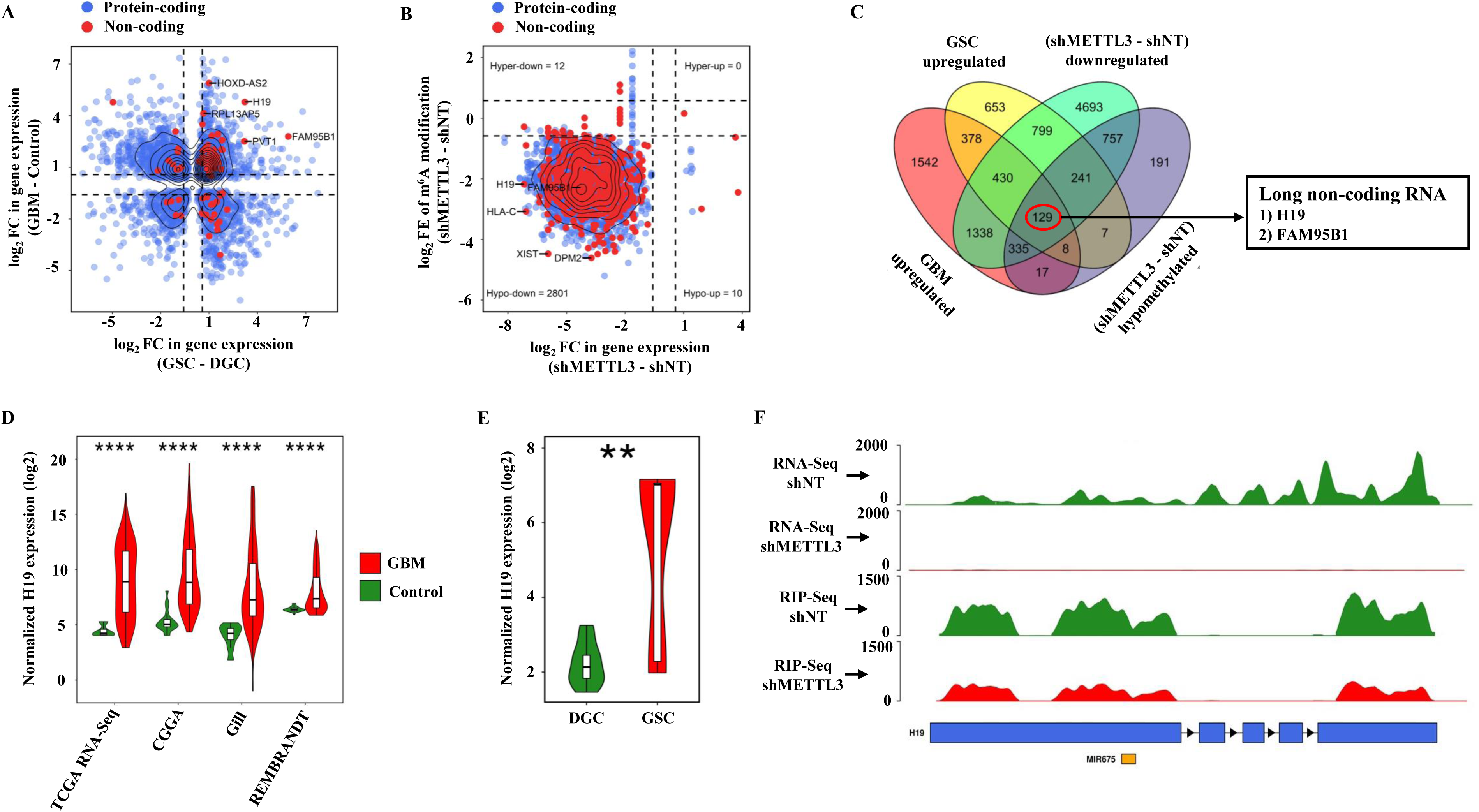
Identification of METTL3 mediated m^6^A modified oncogenic lncRNA. **A.** Scatter plot depicting the pairwise log2 fold change in gene expression between GSC and DGC samples from the GSE54792 cohort (x-axis) and between GBM and control samples from the TCGA RNA-Seq cohort (y-axis). Each point represents a gene, with its coordinates indicating the log_2_FC values in the two cohorts. Non-coding genes substantially upregulated in GBM and GSC are annotated with their respective gene symbols. Quadrant divisions are set at ±0.58 log_2_FC on both axes to identify genes with significant changes in both cohorts. **B.** Scatter plot showing the relationship between log2 fold change in gene expression (x-axis) and log2 fold enrichment of m^6^A peaks (y-axis) from a conjoint analysis of RNA-Seq and m^6^A RIP-Seq data comparing shMETTL3 with shNT conditions. The plot is segmented into four quadrants: hypo m^6^A modified and upregulated genes (lower right), hypo m^6^A modified and downregulated genes (lower left), hyper m^6^A modified and upregulated genes (upper right), and hyper m^6^A modified and downregulated genes (upper left). In the lower-left quadrant, non-coding genes that exhibit substantial hypo m^6^A modification and downregulation in the shMETTL3 condition are labeled with their respective gene symbols. **C.** Four-way Venn diagram illustrating the overlap of coding and non-coding genes across four conditions. The intersections display the total number of genes shared between each subset of conditions. Notably, H19 and FAM95B1 were uniquely identified as the sole non-coding genes common to all four conditions. **D.** Comparative analysis of H19 expression in GBM and control samples across four independent cohorts is illustrated using box-violin plots, which show the gene expression distribution for each group, with GBM samples depicted in red and control samples in green. The y-axis is scaled to represent the log2-transformed normalized gene expression values. Statistical significance between GBM and control groups within each cohort was assessed using the Student’s t-test, with p-values indicated in the panels. **E.** Box-violin plot comparing log2-transformed normalized H19 expression levels between GSC and DGC samples from the GSE54792 cohort. GSC samples are represented in red and DGC samples in green. Statistical significance between GSC and DGC samples was assessed using the Student’s t-test, with p-values indicated in the panel. **F.** Genomic tracks comparing read coverage for H19 between shMETTL3 and shNT conditions in RNA-Seq and m^6^A RIP-Seq experiments. RNA-Seq tracks illustrate changes in transcript abundance while m^6^A RIP-Seq tracks highlight the enrichment or depletion of m^6^A modifications on the H19 transcript, with color-coded tracks distinguishing between shMETTL3 (red) and shNT (green). The y-axis represents read coverage and the x-axis indicates exon boundaries of the H19 gene. Asterisks indicate statistical significance: * for *p* < 0.05, ** for *p* < 0.01, *** for *p* < 0.001, **** for *p* < 0.0001, and ‘ns’ for non-significant differences.

**Figure 2:**
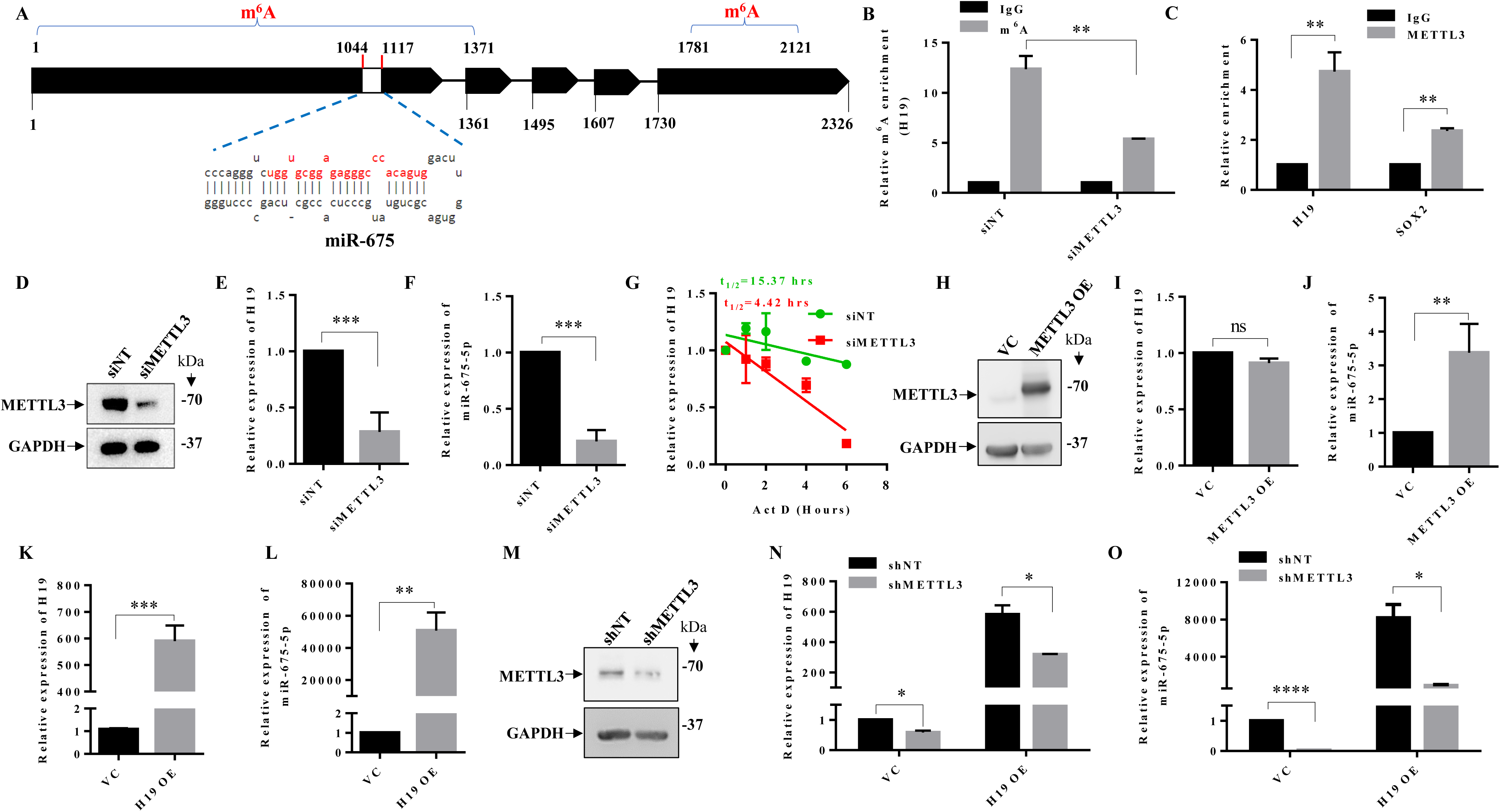
METTL3-mediated m^6^A modification is a critical regulator of H19 stability and miR-675 processing. **A.** The schematic represents the H19 transcript and cognate miR-675 derived from its first exon (the red-colored sequences represent miR-675-5p). **B.** m^6^A pulldown, followed by qRT-PCR measured the enrichment of m^6^A in the H19 transcript. m^6^A enrichment in H19 transcripts in control and METTL3-silencing cells using m6A RIP-qPCR. **C.** The METTL3 binding to the H19 and SOX2 transcripts was quantified by METTL3-RIP-PCR. **D.** METTL3 expression in LN229 cells transfected with control, or METTL3 shRNA was detected by immunoblotting. GAPDH served as the control. **E & F.** The expression of H19 and miR-675-5p were measured by qRT-PCR in METTL3 silenced LN229 cells. **G.** The stability of the H19 transcript was measured at indicated time points post Actinomycin D (5 μg/ml) treatment in shNT and shMETTL3-transfected LN229 cells by qRT-PCR. The log2 ratio of the remaining H19 was plotted using linear regression after normalizing to the 0th hour of the respective condition. **H.** METTL3 expression in METTL3 overexpressed LN229 cells was detected by immunoblotting. GAPDH served as the control. **I & J.** The expression of H19 and miR-675-5p were measured by qRT-PCR in METTL3 overexpressing LN229 cells. **K & L.** The expression levels of H19 and miR-675-5p were measured by qRT-PCR in H19-overexpressing LN229 cells. **M.** The expression of METTL3 were measured by immunoblotting in shNT and shMETTL3 LN229 cells. **N & O.** H19 and miR-675-5p were measured in stable H19 overexpressed with METTL3 silenced cells. Data are shown as mean ± SD (n=3). Asterisks indicate statistical significance: * for *p* < 0.05, ** for *p* < 0.01, *** for *p* < 0.001, **** for *p* < 0.0001, and ‘ns’ for non-significant differences.

Based on m^6^A RIP-Seq, we found that H19 is m^6^A-modified with two m^6^A peaks in the first and the fifth exons (**Figure 2A**; (Visvanathan et al., 2019b)). Validation experiments demonstrate a significant enrichment of H19 transcripts in anti-m^6^A immunoprecipitation (IP) with a significant reduction in METTL3 silenced glioma cells, suggesting that H19 undergoes METTL3-dependent m^6^A modification (**Figure 2B**). An anti-METTL3 IP also showed significant enrichment of H19 (**Figure 2C**). LN229 glioma cells with METTL3 silencing showed a substantial reduction in H19 and miR-675 levels (**Figure 2D**, **E, and F**). Actinomycin-treated METTL3 silenced LN229 glioma cells showed a significant decrease in the half-life of H19 (4.20 hrs in siMETTL3 over long hours, more than 6 hrs in siNT conditions), suggesting that METTL3-dependent m^6^A modification promotes H19 stability (**Figure 2G**). Interestingly, exogenous overexpression of METTL3 in LN229 glioma cells (**Figure 2H**) resulted in a several-fold increase in miR-675 levels with no change in H19 levels (**Figure 2I and J**), indicating the existence of differential processing of m^6^A-modified H19 lncRNA concerning H19 stability vs. miRNA processing. In contrast, exogenous overexpression of H19 in LN229 glioma cells increased H19 and miR-675 levels by several folds (**Figures 2K and L**). However, exogenous overexpression of H19 in METTL3-silenced cells (**Figure 2M**) showed a significant difference in the elevation of H19 and miR-675 levels. While the absence of METTL3 affected the expression of both H19 and miR-675 expressed from exogenously expressed H19, miR-675 expression was severely affected (**88.7%)** compared to H19 (**45.3 %)** levels (**Figure 2N**, **and O**), adding more strength to the earlier conclusion that m^6^A modification regulates H19 stability and miR-675 processing differently.

### H19 transcript processing is controlled by m^6^A reader proteins - HNRNPA2B1 and IGF2BP2

The fate of m^6^A-modified RNAs is determined by m^6^A reader proteins (Visvanathan & Somasundaram, 2018). To dissect the processing of m^6^A-modified H19 transcript, we explored a family of RNA binding proteins (RBPs) that are potential m^6^A readers. We intersected the RBPs having a positive correlation with H19 at transcript levels in GBM transcriptome data (**Supplementary Table 8**) with H19-bound RBPs that are conserved in human (**Supplementary Table 9)** and mouse (**Supplementary Table 10**) curated from the ENCORI database (https://rnasysu.com/encori/index.php). This analysis identified two RBPs, HNRNPA2B1 and IGF2BP2, as potential H19 m^6^A reader proteins (**Figure 3A**). H19 transcript levels show a significant positive correlation with IGF2BP2 and HNRNPA2B1 transcript levels in GBM (**Figure 3B and C**). We found a transcript upregulation of HNRNPA2B1 and IGF2BP2 in multiple GBM data sets compared to normal brain samples (**Supplementary Figure 2A and B**). To confirm the interaction between H19 and selected RBPs, we tested the ability of streptavidin-binding RNA aptamer fused H19 (**Figure 3D**; H19-4S1m) transcript (Xu et al., 2021)to bind to the RBPs. Streptavidin beads pulldown of H19-4S1m efficiently brought down H19 transcript (**Figure 3E)** and HNRNPA2B1/IGF2BP2 proteins (**Figure 3F**) compared to VC-4S1m pulldown. In a reverse experiment, an RNA immunoprecipitation (RIP) experiment using antibodies against HNRNPA2B1 or IGF2BP2 efficiently brought down both RBPs (**Figure 3G)** and H19 transcript (**Figure 3H**). To study the requirement of the chosen RBPs on H19 stability and miR-675 processing, we studied the impact of RBP silencing on H19 processing. HNRNPA2B1 or IGF2BP2 silencing decreased the H19 and miR-675 levels (**Figure 3I**, **J, and K**). Actinomycin experiment showed a reduced half-life for H19 transcript in HNRNPA2B1 or IGF2BP2 silenced glioma cells (**Figure 3L**), suggesting that HNRNPA2B1 or IGF2BP2 binding promotes stability of H19 transcript.

**Figure 3:**
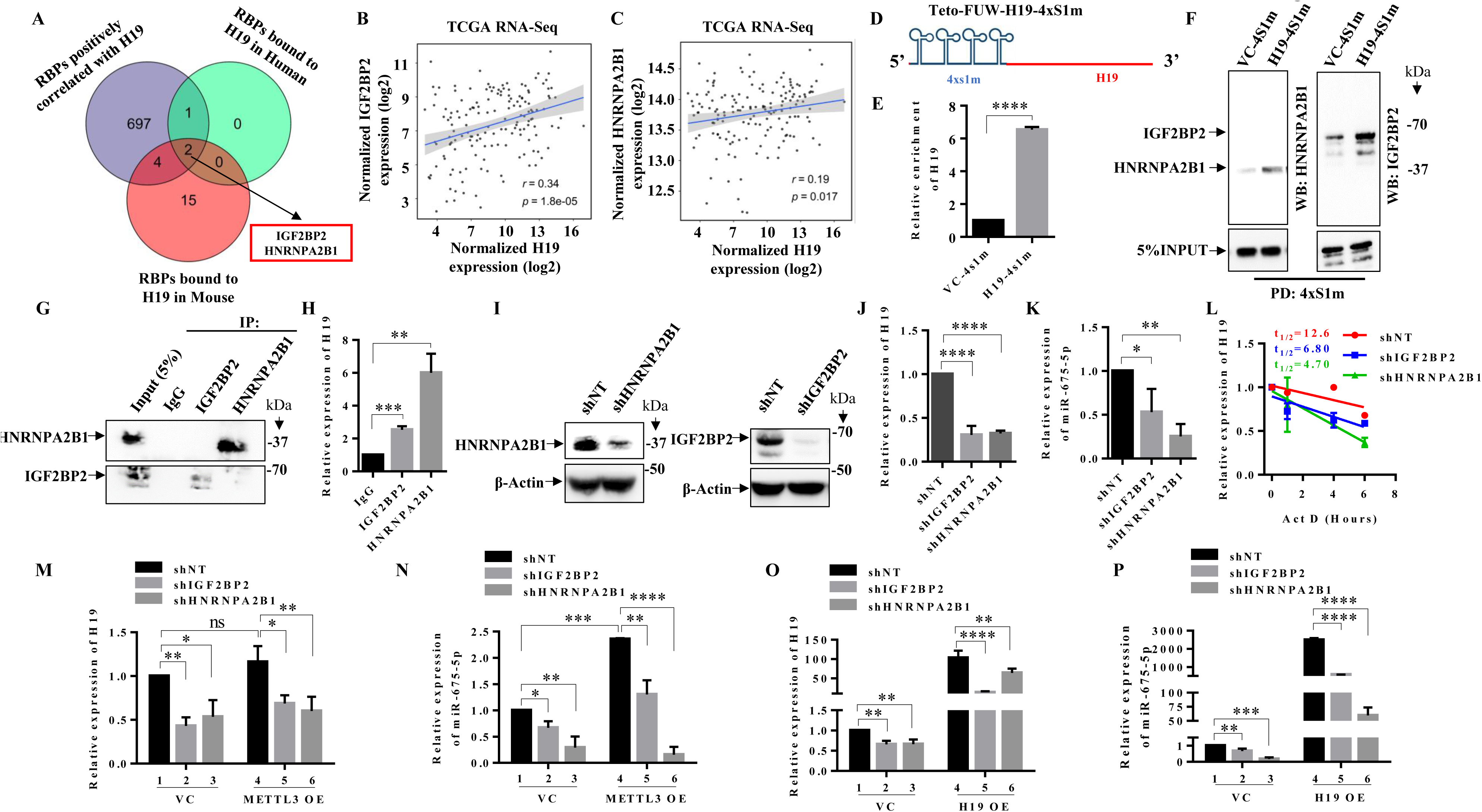
H19 transcript processing is controlled by m^6^A reader proteins HNRNPA2B1 and IGF2BP2. **A.** The Venn diagram depicts the number of RBPs that are (1) positively correlated with H19 in the TCGA RNA-Seq cohort, (2) experimentally validated to bind H19 in humans, and (3) experimentally validated to bind H19 in mice, as obtained from the ENCORI database. HNRNPA2B1 and IGF2BP2 are highlighted as the only two RBPs that bind to H19 and are conserved in humans and mice. **B.** The scatter plot shows log2-transformed H19 expression on the x-axis and IGF2BP2 expression on the y-axis. The regression line with a 95% confidence interval indicates the overall correlation trend, with Pearson’s correlation coefficient (r) and p-value (p) provided to quantify the strength and statistical significance of the correlation. **C.** The scatter plot shows log2-transformed H19 expression on the x-axis and HNRNPA2B1 expression on the y-axis. The regression line with a 95% confidence interval indicates the overall correlation trend, with Pearson’s correlation coefficient (r) and p-value (p) provided to quantify the strength and statistical significance of the correlation. **D.** The schematic represents 4s1m-H19 transcript structure for pulldown to validate the interaction between H19 and reader proteins. **E.** We performed a pulldown of 4s1m-H19 using streptavidin magnetic beads, followed by quantification of H19 using qRT-PCR. **F.** We conducted the pulldown of 4s1m-H19, followed by protein elution to demonstrate the interaction between H19 and reader proteins (HNRNPA2B1 and IGF2BP2) using immunoblotting. **G.** We conducted immunoprecipitation using antibodies against IGF2BP2 and HNRNPA2B1, and the eluted proteins were subsequently subjected to SDS-PAGE, followed by immunoblotting using antibodies against HNRNPA2B1 and IGF2BP2. **H.** We performed immunoprecipitation using antibodies against IGF2BP2 and HNRNPA2B1, and the eluted RNA was subsequently analyzed using qRT-PCR. **I.** The protein levels of HNRNPA2B1 and IGF2BP2 were assessed by immunoblotting in HNRNPA2B1 and IGF2BP2 silenced LN229 cells compared to control cells. **J & K.** We conducted qRT-PCR for H19 and miR-675 in HNRNPA2B1 and IGF2BP2 silenced LN229 cells. **L.** The stability of the H19 transcript was measured at indicated time points post Actinomycin D (5 μg/ml) treatment in shHNRNPA2B1 and shIGF2BP2-transfected LN229 cells by qRT-PCR. The log2 ratio of the remaining H19 was plotted using linear regression after normalizing to the 0th hour of the respective condition. **M & N.** H19 and miR-675-5p levels were assessed via qRT-PCR in METTL3 overexpressed cells where HNRNPA2B1 and IGF2BP2 were silenced. **O & P.** H19 and miR-675-5p levels were assessed via qRT-PCR in H19 overexpressed LN229 cells where HNRNPA2B1 and IGF2BP2 were silenced. Data are shown as mean ± SD (n=3). Asterisks indicate statistical significance: * for *p* < 0.05, ** for *p* < 0.01, *** for *p* < 0.001, **** for *p* < 0.0001, and ‘ns’ for non-significant differences.

To delineate the functions of these two RBPs in H19 processing, we analyzed the impact of exogenous overexpression of METTL3 in RBP-silenced conditions. We showed above that the exogenous overexpression of METTL3 promoted miR-675 processing with no change in H19 transcript levels (**Figure 2I and J**). To elucidate the requirements of the two RBPs on the endogenous H19 processing, we tested the impact of METTL3 overexpression on endogenous H19 processing in either of the RBPs silenced glioma cells. As expected, exogenous overexpression of METTL3 increased the miR-675 levels (**Figure 3N**; **compare bar 4 with 1**) with no change in H19 transcript levels (**Figure 3M**; **compare bar 4 with 1**). In HNRNPA2B1 or IGF2BP2 silenced conditions, a significant reduction was seen in the H19 transcript levels (**Figure 3M**; **compare bars 2 and 3 with 1**). Interestingly, the reduction seen in H19 transcript levels in HNRNPA2B1 or IGF2BP2 silenced cells was not altered upon exogenous overexpression of METTL3 (**Figure 3M**; **compare bars 5 and 6 with 4**), adding strength to the fact that RBPs, as m^6^A reader proteins, function downstream to METTL3. In contrast, we observed a differential requirement of RBPs under METTL3 overexpressed conditions for miR-675 processing. In METTL3 overexpressing conditions, miR-675 level was reduced only by 44.7 % in IGF2BP2 silenced conditions (**Figure 3N**; **compare bar 5 with 4**), which could be explained by the reduced H19 levels in IGF2BP2 silenced conditions by about 50% (Figure 3M; compare bar 5 with 4). In contrast, the miR-675 level was reduced drastically in HNRNPA2B1 silenced conditions (93.4%) compared (**Figure 3N**; **compare bars 6 with 4),** suggesting that HNRNPA2B1 plays a major role in miR-675 processing. As a control experiment, we show that transcript levels of IGF2BP2 and HNRNPA2B1 are not affected in glioma cells with METTL3 silencing or METTL3 overexpression (**Supplementary Figure 2C and D**).

Next, we tested the importance of RBPs on the processing of H19 expressed from an exogenously introduced H19 construct (H19-4S1m) by using RBP silenced cells. We showed above that exogenous overexpression of H19 increased both H19 transcript levels and miR-675 levels (**Figure 2K and L**). As expected, the exogenous overexpression of H19 resulted in a several-fold increase in H19 transcript levels (**Figure 3O**; **compare bar 4 with 1**) and miR-675 levels (**Figure 3P**; **compare bar 4 with 1**). The increase in H19 transcripts from exogenously overexpressed H19 is compromised severely in IGF2BP2 silenced cells (**87%; Figure 3O**; **compare bar 5 with 4**) compared to HNRNPA2B1 silenced cells (**38%; Figure 3O**; **compare bar 6 with 4),** suggesting that IGF2BP2 may play a major role in H19 stability. In contrast, the increase in miR-675 levels from exogenously overexpressed H19 is compromised in HNRNPA2B1 silenced cells (**97.6%**; **Figure 3P**; **compare bar 6 with 4)** compared to IGF2BP2 silenced cells (**77%; Figure 3Q**; **compare bar 5 with 4**), suggesting that HNRNPA2B1 may play a major role in miR-675 processing. From these results, we conclude that m^6^A reader proteins play specific roles in H19 processing downstream to m^6^A modification, with IGF2BP2 playing an important role in H19 stability and HNRNPA2B1 playing an important role in miR-675 processing.

### Identification of m^6^A modification sites and their role in H19 processing

To investigate the differential role played by m^6^A modification on the processing of H19, we resorted to identifying the m^6^A modification sites on the H19 transcript. m^6^A-RIP sequencing identified that there are two m^6^A peaks with a peak score of 2.174 located between nucleotides 1 to 1371, corresponding to exon 1, and 0.281 between nucleotides 1781 to 2121, corresponding to exon 5 (**Figure 2A**; (Visvanathan et al., 2019b)). To locate the m^6^A sites precisely, we subjected the H19 transcript nucleotide sequence to SRAMP (**s**equence-based **R**NA **a**denosine **m**ethylation site **p**redictor) prediction tool (https://www.cuilab.cn/sramp). SRAMP predicted a total of eight m^6^A sites, with six sites in the first exon and one site in the last exon (**Figure 4A**). For further studies, we took the two high-confidence m^6^A sites predicted in the first exon; site #2 is located on the loop region of miR-675 locus in the H19 transcript, and site #1 is located at 311 nucleotides upstream of site #2. To study the contribution of these two m^6^A sites, we introduced mutations in each of the sites (**A to G; Figure 4A**) into H19-4S1m construct to create H19 mut#1-4S1m and H19 mut#2-4S1m constructs. The WT and mutant H19 transcripts were purified from H19-4S1m, H19 mut#1-4S1m, and H19 mut#2-4S1m transfected cells and incubated with anti-m^6^A antibodies. The bound antibody was brought down using the streptavidin pull-down, and the extent of m^6^A modification in either WT or mutant RNAs was measured by quantifying the anti-m^6^A antibody binding by a western blot with a secondary antibody. The extent of anti-m^6^A binding is reduced by approximately 50% in either of the mutants (**Figure 4B**), indicating the fact that either of the mutations has affected the m^6^A modification partially, with the possibility that in each of the mutants, the remaining WT site is probably modified. Next, to study the relative roles of the two m^6^A sites in H19 processing, we transfected H19-4S1m, H19 mut#1-4S1m, and H19 mut#2-4S1m constructs into HEK293 cells, which express undetectable endogenous H19 transcripts, and measured the H19 and miR-675 levels derived from the parent transcripts. While we observed a significant reduction in H19 transcript and miR-675 levels in either of the mutant H19 construct transfected cells, we found some interesting differential impact by m^6^A site #1 vs 2 mutations. While the H19 level was most affected in H19 mut #2-4S1m transfected cells (**Figure 4C**; **compare bars 4 with 2)**, the miR-675 levels were most affected in H19 mut #1-4S1m transfected cells (**Figure 4D**; **compare bars 3 with bar 2**). This result suggests that m^6^A site #2 may have a larger role in H19 stability, while miR-675 processing is mainly controlled by m^6^A site #1. Next, we tried to integrate the unique requirement of specific m^6^A sites versus RBPs in H19 processing. Our previous results, as shown above, identified that H19 stability is mainly controlled by IGF2BP2, while the miR-675 processing is controlled by HNRNPA2B1 (**Figure 3M**, **N, O, and P**). We transfected the WT and mutant H19 constructs in IGF2BP2 or HNRNPA2B1 silenced cells and analyzed the impact of specific m^6^A sites and RBPs on H19 processing. As expected, H19 levels expressed from the WT H19 (H19-4S1m) construct are reduced significantly in IGF2BP2 (**Figure 4E**; **compare bar 2 with 1**) silenced cells. However, the level of H19 transcript derived from H19 mut #2-4S1m construct, unlike H19 mut #1-4S1m construct, failed to show any reduction in IGF2BP2 silenced cells, suggesting that m^6^A site # 2 may mediate IGF2BP2 binding to promote H19 stability. In another experiment, we found that miR-675 expressed from the WT H19 construct is reduced significantly in HNRNPA2B1 (**Figure 4F**; **compare bar 2 with 1**) silenced cells. However, the level of miR-675 derived from the H19 mut #1-4S1m construct, unlike the H19 mut #2-4S1m construct, failed to show any further reduction (**Figure 4F**; **compare bar 4 with 3**), suggesting that m^6^A site #1 may mediate HNRNPA2B1 binding to promote miR-675 processing. Thus, we conclude from these results that m^6^A reader proteins, IGF2BP2 and HNRNPA2B1, bind to specific m^6^A sites to regulate H19 processing differentially.

**Figure 4:**
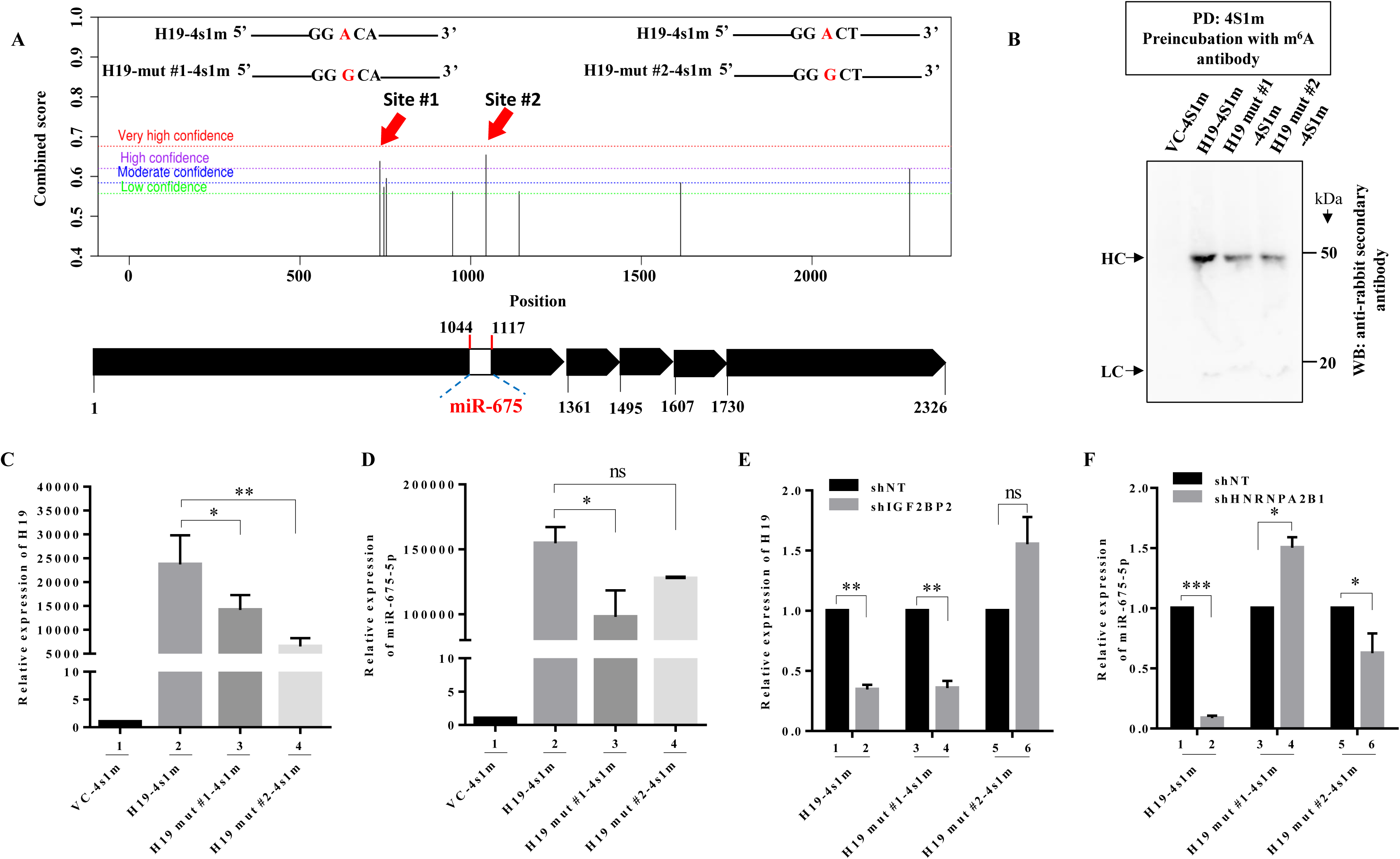
Identification of m^6^A modification sites and their role in H19 processing. **A.** The schematic represents the prediction score distribution for the m^6^A motif along the H19 sequence using SRAMP. The bottom part of the diagram shows the intronless H19 transcript containing the miR-675 locus, aligned with the m^6^A motif distribution. **B.** We have demonstrated that m^6^A modification in H19-4s1m, H19 mut#1-4s1m, and H19 mut#2-4s1m transcripts through streptavidin magnetic beads mediated 4s1m pulldown followed by m^6^A antibody incubation, and the eluted bound antibody was detected by immunoblotting with mouse-secondary antibody. **C & D.** H19 and miR-675-5p expressions were measured using qRT-PCR in H19-4s1m, H19 mut#1-4s1m, and H19 mut#2 expressed HEK293T cells. **E & F.** We investigated the expression of H19 and miR-675-5p in cells expressing H19-4s1m, H19 mut#1-4s1m, and H19 mut#2 while IGF2BP2 and HNRNPA2B1 were silenced. Data are shown as mean ± SD (n=3). Significance is denoted by asterisks: * for p < 0.05, ** for p < 0.01, *** for p < 0.001, **** for p < 0.0001, and ‘ns’ for non-significant differences.

### H19-METTL3-IGF2BP2-HNRNPA2B1-miR-675-CALN1 axis regulates glioma migration and imparts a poor prognosis

The function of H19 in glioma progression and its targets has been extensively studied (Chen et al., 2018; Fazi et al., 2018; Jiang et al., 2016; W. Li et al., 2016; P. Liu et al., 2021; G. Wang et al., 2022; W. Wu et al., 2017; Zhou et al., 2022). However, the role of miR-675 and its targets has not been thoroughly investigated. To understand the importance of the H19-METTL3 axis, we have explored miR-675 targets and their impact on glioma progression. To identify potential targets of miR-675 in glioma development, we used miRTarBase (https://mirtarbase.cuhk.edu.cn/~miRTarBase/miRTarBase_2022/php/index.php). Among the predicted targets (**Supplementary Table 11**), we chose to work on Calneuron 1 (CALN1) for the following reasons. CALN1 encodes a calcium-binding protein and is implicated in signal transduction (Y. Q. Wu et al., 2001). CALN1 has already been demonstrated as the target of miR-675 target in osteosarcoma and gastric cancer (Gong et al., 2018; H. Li et al., 2014). Pan cancer data analysis revealed that CALN1 is specifically downregulated in GBM compared to control brain samples (**Figure 5A**). The GBM-specific downregulation of CALN1 is validated in multiple GBM transcriptome data sets (**Figure 5B**). The analysis of the Genotype-Tissue Expression (GTEx) of normal tissues revealed that CALN1 is highly expressed in normal brain tissue, while H19 is undetectable in normal brain tissue samples (**Supplementary Figure 3A and B**). Silencing METTL3 increased the CALN1 transcript levels (**Figure 5C**). While silencing either IGF2BP2 or HNRNPA2B1 increased the CALN1 transcript levels, the impact of HNRNPA2B1 loss on the CALN1 level increase was much higher (**Figure 5D**). METTL3 silencing reduced miR-675 levels and can be rescued by exogenous overexpression of miR-675 mimic (**Figure 5E**). METTL3 silencing decreased glioma cell migration and invasion, which could be reversed by exogenous overexpression of miR-675 mimic (**Figure 5F and G**; **Supplementary Figure 4A, B, C, and D**). Similarly, silencing of either IGF2BP2 or HNRNPA2B1 decreased glioma cell migration, which could be reversed by exogenous overexpression of miR-675 mimic (**Figure H; Supplementary Figure 4E, F, and G**).

**Figure 5:**
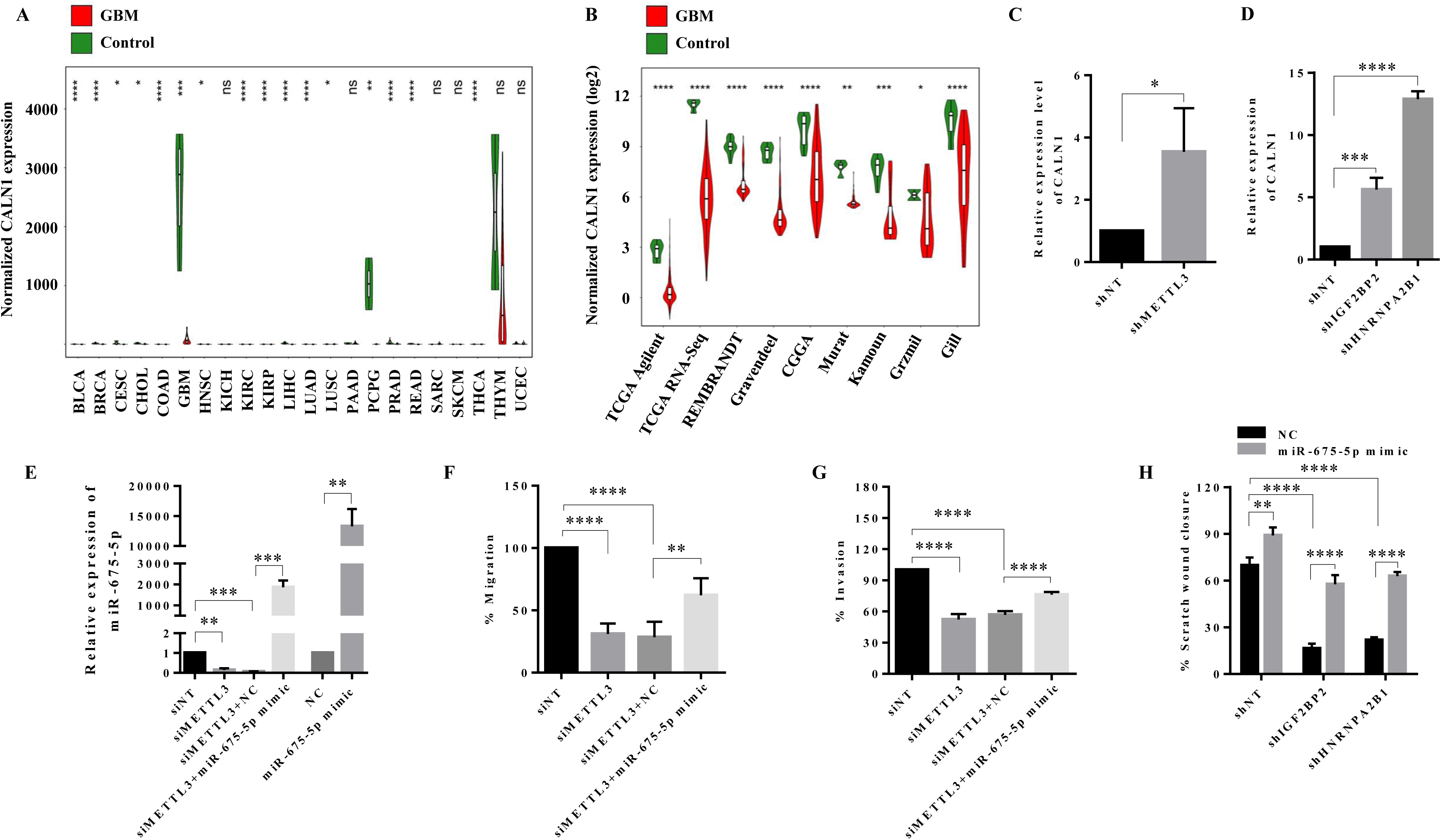
METTL3-H19-IGF2BP2-HNRNPA2B1-miR-675 axis inhibits CALN1 to promote glioma migration. **A.** This figure shows box-violin plots comparing CALN1 expression levels in GBM (red) and control (green) samples across 22 cancer types obtained from the TCGA Pan-Cancer Atlas Project. The y-axis displays RSEM-normalized gene expression values. Statistical differences between GBM and control groups were assessed with the Student’s t-test, and p-values are shown in the panels. **B.** Comparative analysis of CALN1 expression in GBM and control samples across nine independent cohorts. The box-violin plots illustrate the gene expression distribution for each group within each cohort, with GBM samples depicted in red and control samples in green. The y-axis is scaled to represent the log2-transformed normalized gene expression values. Statistical significance between GBM and control groups within each cohort was assessed using the Student’s t-test, with p-values indicated in the panels. **C & D.** CALN1 expression was quantified using qRT-PCR in shMETTL3, shHNRNPA2B1, and shIGF2BP2 cells compared to the control. **E.** LN229 cells with silenced METTL3 were transfected with either a miR-675-5p mimic or negative control (NC), followed by qRT-PCR analysis to measure miR-675-5p expression. **F & G.** The Boyden chamber invasion and migration assays were quantified on LN229 cells with silenced METTL3 and a miR-675 mimic. **H.** The migratory capacity of cells with silenced IGF2BP2 and HNRNPA2B1 transfected with a miR-675-5p mimic was evaluated using a scratch wound assay. Data are shown as mean ± SD (n=3). Significance is denoted by asterisks: * for p < 0.05, ** for p < 0.01, *** for p < 0.001, **** for p < 0.0001, and ‘ns’ for non-significant differences.

To further explore the clinical relevance of these findings, we investigated the GBM transcriptome data. Correlation analysis revealed a significant negative correlation of CALN1 transcript levels with H19, METTL3, IGF2BP2, and HNRNPA2B1 transcripts (**Supplementary Figure 3C**). We also found a significant positive correlation between H19, METTL3, IGF2BP2, and HNRNPA2B1 transcripts (**Supplementary Figure 3C**). Survival analysis in GBM patients provided significant evidence that the H19-METTL3-HNRNPA2B1-miR675-CALN1 axis indeed is operating in glioma (**Figure 6A**). In addition, low CALN1 transcript levels predicted a poor prognosis (**Figure 6B**; **Supplementary Figure 3D**). CALN1 low/H19 high transcript levels predicted a worse prognosis (**Figure 6C**; **Supplementary Figure 3E**). In the CGGA data set, CALN1 low/H19 high GBMs with high levels of METTL3 or HNRNPA2B1 predicted a worse prognosis (**Figure 6D and E**). However, IGF2BP2 transcript levels failed to stratify the CALN1 low/H19 high GBMs with a significant survival difference (**Figure 6F**). Further, CALN1 low/H19 high/METTL3 high GBM patients with high levels of HNRNPA2B1 predicted a worse prognosis (**Figure 6G**). However, IGF2BP2 transcript levels failed to stratify the CALN1 low/H19 high/METTL3 high GBMs with a significant survival difference (**Figure 6H**). The ability of HNRNPA2B1 but not the IGF2BP2 transcript levels to predict survival in the context of H19/METTL3/miR-675/CALN1 axis reflects the fact that HNRNPA2B1 plays a major role in miR-675 processing of m^6^A modified H19 transcripts. From these results, we conclude that CALN1, a bonafide target of miR-675, is regulated by the H19-METTL3-HNRNPA2B1-miR-675 axis in glioma to promote migration and invasion.

**Figure 6:**
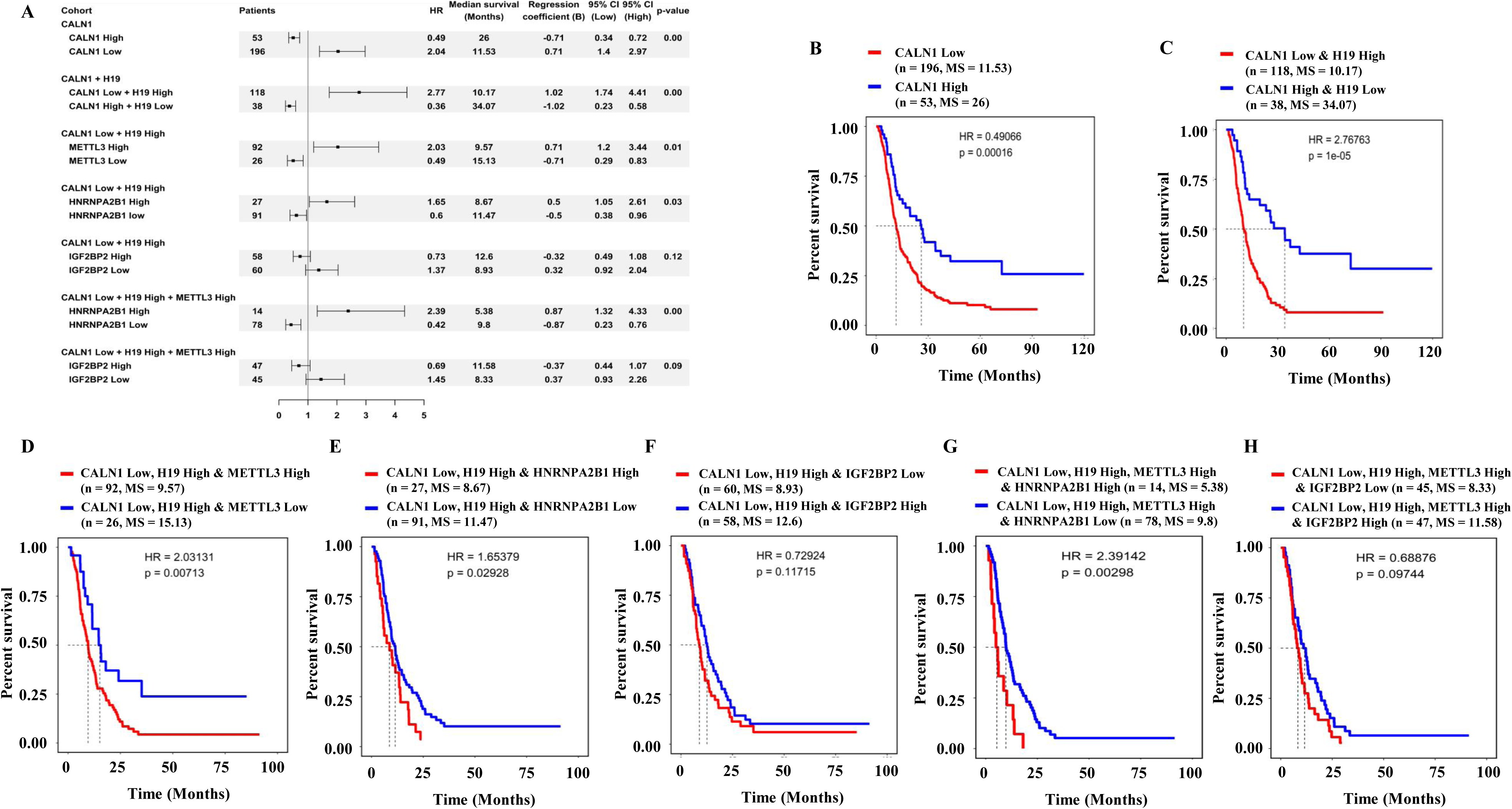
METTL3-H19-IGF2BP2-HNRNPA2B1-CALN1 axis is associated with glioma progression. **A.** The forest plot summarizes the hazard ratios for survival outcomes associated with high or low expression of individual genes or gene combinations in the CGGA cohort. Hazard ratios of individual studies are shown as squares with horizontal lines representing their confidence intervals. The vertical line denotes no effect (i.e. hazard ratio of 1). **B-H.** Kaplan-Meier survival curves comparing overall survival probabilities between patients with low versus high expression of individual genes or gene combinations in the CGGA cohort. Survival time in months is shown on the x-axis, and overall survival probability is shown on the y-axis. The red line represents the patient group associated with a worse prognosis, while the blue line corresponds to the group with a better survival outcome. The figure includes the hazard ratio (HR) and p-value for assessing survival differences, as well as the number of patients (n) and median survival in months (MS) for each group. Significance is denoted by asterisks: * for p < 0.05, ** for p < 0.01, *** for p < 0.001, **** for p < 0.0001, and ‘ns’ for non-significant differences.

## Discussion

H19 and the cognate miR-675 are derived from an lnc-pri-miRNA locus. H19 has been to have both miR-675 dependent and -independent functions (Keniry et al., 2012). While the H19/miR-675 axis has been shown to promote glioma development, the precise mechanism behind H19 processing, particularly the H19 stability versus miR-675 formation, is unknown. In this study, we demonstrate that H19 undergoes METTL3-dependent m^6^A modification which is essential for its stability and processing to form miR-675. We also found that m^6^A modification regulates H19 processing with m^6^A reader proteins, IGF2BP2 and HNRNPA2B1 regulating the H19 stability and miR-675 processing respectively. Finally, we found that CALN1, a bonafide target of miR-675 (Gong et al., 2018; H. Li et al., 2014), is under the control of H19/METTL3/IGF2BP2/HNRNPA2B1/miR-675 axis with GBMs having low CALN1/high H19 transcript levels predicting worse prognosis.

We found that H19 is expressed at a higher level in GBM and low-grade glioma. The patient-derived GSCs also express high levels of H19 compared to DGCs. Our results also showed that H19 undergoes METTL3-dependent m^6^A modification as the m^6^A peak was lost in METTL3-silenced cells. We also found that H19 stability requires m^6^A modification; the H19 transcript levels without m^6^A modification in METTL3 silenced cells are reduced drastically. Interestingly, while METTL3 overexpression significantly increased miR-675 levels, it failed to alter H19 levels, indicating that m^6^A modification selectively influences miR-675 processing, perhaps using a different mechanism compared to H19 stability. METTL3-dependent m^6^A modification appears to stabilize H19 but is more crucial for miR-675 processing, suggesting a nuanced regulatory mechanism where m^6^A modifications influence distinct aspects of H19’s functionality.

We found that two RBPs, IGF2BP2 and HNRNPA2B1, which are m^6^A reader proteins (Alarcón et al., 2015; Huang et al., 2018), interact with H19, and their absence reduced the H19 and miR-675 levels. The m^6^A reader proteins appear to play distinct roles in H19 stability and miR-675 processing. Silencing these proteins results in decreased levels of both H19 and miR-675. Silencing with HNRNPA2B1 primarily affects miR-675 processing, and IGF2BP2 mainly influences H19 stability. This differentiation underscores the complexity of m^6^A-mediated regulation and the specificity of m^6^A reader proteins in modulating lncRNA functions.

The identification of specific m^6^A modification sites within the H19 transcript further elucidates its regulatory mechanisms. Two high-confidence m^6^A sites were identified in the first exon, with site-specific mutations demonstrating differential impacts on H19 stability and miR-675 processing. Mutation at m^6^A site #2 predominantly affected H19 stability, while mutation at site #1 significantly impacted miR-675 processing, reinforcing the idea that distinct m^6^A sites have specific regulatory roles. Further, the mutation in the m^6^A site #2 failed to decrease the H19 levels in IGF2BP2 silenced cells, suggesting that the m^6^A site #2 may promote IGF2BP2 binding to stabilize the H19 transcript. In contrast, a mutation in the m^6^A site #1 failed to decrease miR-675 levels in HNRNPA2B1 silenced cells, suggesting that site #1 may promote HNRNPA2B1 interaction to augment miR-675 formation. Thus, we propose a mechanistic model for H19 processing (**Figure 7**), which proposes that m^6^A modification promotes IGF2BP2 and HNRNPA2B1 binding to H19 transcript at specific sites to promote H19 stability versus miR-675 formation, respectively.

**Figure 7:**
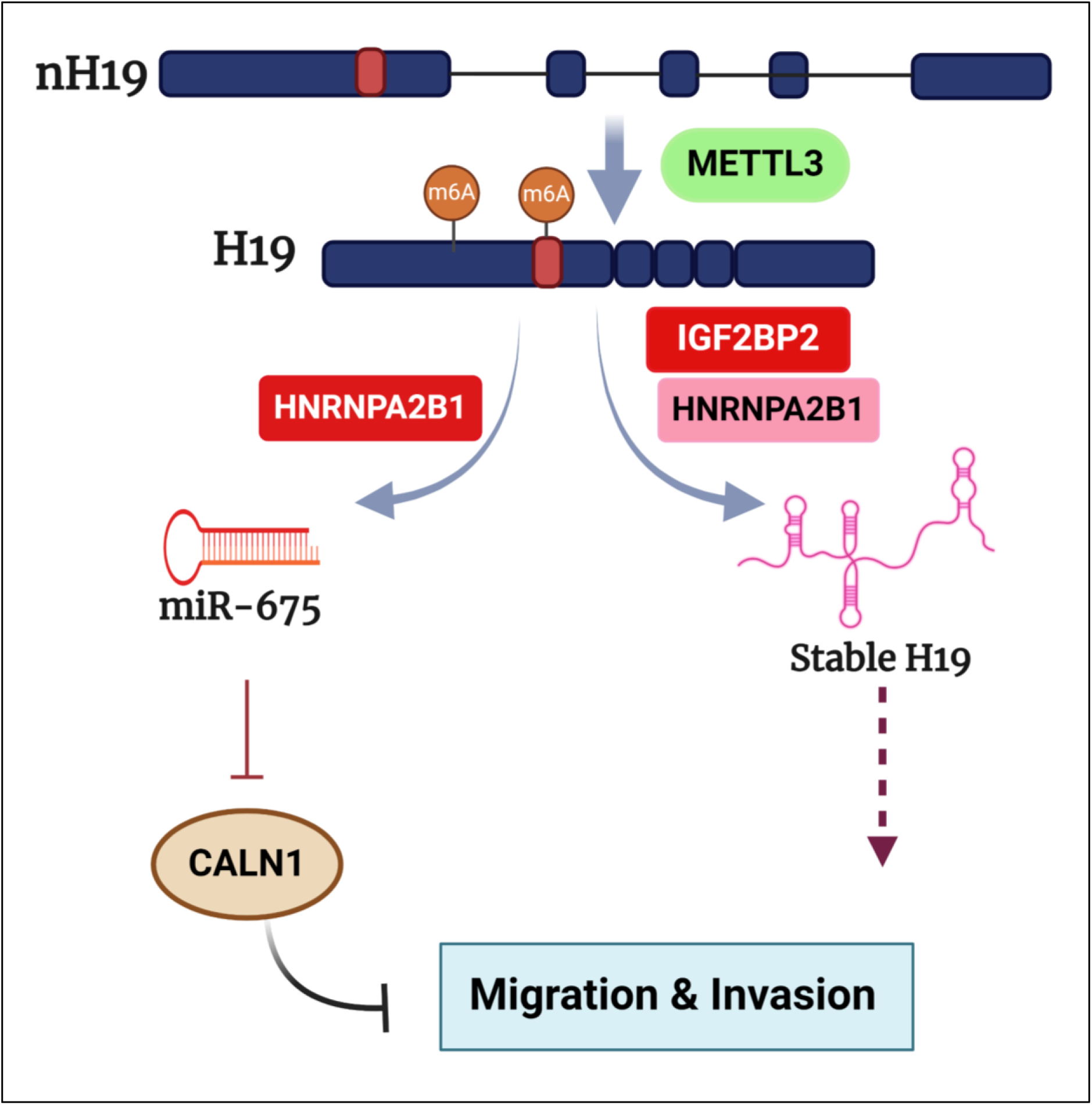
Proposed working model of this study. Decoding the role of METTL3 mediated m^6^A modification on H19 (created with BioRender.com).

The study also found that the H19-METTL3-IGF2BP2-HNRNPA2B1-miR-675 regulatory axis promotes glioma cell migration and invasion by downregulating Calneuron 1 (CALN1), a target of miR-675. Silencing METTL3, IGF2BP2, or HNRNPA2B1 increases CALN1 levels, reducing glioma cell migration and invasion. These effects are reversible with miR-675 overexpression, highlighting the axis’s critical role in glioma pathogenesis. While low levels of CALN1 predicted a poor prognosis, low CALN1/high H19 levels predicted a worse prognosis. Further, high METTL3 in the low CALN1/high H19 group continues to predict a worse prognosis. Interestingly, higher levels of HNRNPA2B1, but not IGF2BP2, predicted worse prognosis in both low CALN1/high H19 and low CALN1/high H19/high METTL3 GBMs indicating the fact that HNRNPA2B1 levels play a major role in H19/METTL3/miR675/CALN1 axis.

The findings underscore the importance of m^6^A modification in regulating lncRNA functions, particularly H19, in glioma progression. METTL3 and m^6^A reader proteins differentially regulate H19 stability and miR-675 processing, which in turn influences glioma cell migration and invasion through the downregulation of CALN1. This complex regulatory network offers potential therapeutic targets for GBM, emphasizing the need for further research into m^6^A modifications and their broader implications in cancer biology.

### Author contributions

SJ planned and carried out all the experiments; AC conducted all bioinformatic analyses; KS prepared the manuscript, planned the study, and led the project.

## Acknowledgments

The results published here are, in whole or part, based upon data generated by The Cancer Genome Atlas pilot project established by the NCI and NHGRI. Information about TCGA and the investigators and institutions that constitute the TCGA research network can be found at http://cancergenome.nih.gov/. We acknowledge the shRNA consortium (Prof. Subba Rao), IISc, India, for shRNA constructs. We acknowledge Prof. Dung-Fang Lee’s lab (University of Texas, USA) for providing the 4s1m-H19 construct. SJ acknowledges NPDF, DBT RA, and DST for fellowship. For research grants, KS acknowledges CEFIPRA, DBT, DST, ICMR, and CSIR (Govt. of India). Infrastructure supported by DST FIST, DBT-IISc partnership program, and UGC is acknowledged. KS is awarded the J. C. Bose Fellowship from DST.

